# Learning Allosteric Interactions in Gα Proteins from Molecular Dynamics Simulations

**DOI:** 10.1101/2024.10.31.621204

**Authors:** Yiping Yu, Maohua Yang, Wenning Wang

## Abstract

Gα is a key subunit of heterotrimeric guanine-nucleotide-binding regulatory proteins, yet its conformational dynamics are not fully understood. In this study, we developed a Transformer-based graph neural network framework, Dynamic-Mixed Transformer (DMFormer), to investigate conformational dynamics of Gαo. DMFormer achieved an AUC of 0.75 on the training set, demonstrating robustness in distinguishing active and inactive states. The interpretability of the model was enhanced using integrated gradients, identifying the Switch II as a critical motif in stabilizing the active state and revealing distinct movement patterns between GTPase and α-Helix domains. Our findings suggest that the conformational rigidity of the Q205L mutant in the Switch II segment leads to persistent activation. Overall, our study showcases DMFormer as an effective tool for analyzing protein conformational dynamics, offering valuable insights into activation mechanisms of Gα protein.

## INTRODUCTION

Molecular dynamics (MD) simulations have become a widely used computational tool due to their ability to provide fine-grained dynamic information. However, they come with high computational costs, especially when observing processes occurring over long time scales. Integrated variables, such as free energy and contact maps, are used to quantify protein motion and are typically obtained from complete ensembles. One of the main challenges in MD simulations over long time scales is the adequate sampling of rare or transition states, which are characterized by high energies and low probabilities.

Collective variables (CVs) have provided advanced insights into MD trajectories and can be chosen based on prior knowledge, such as ϕ and ψ dihedral angles, distances between key residues. CVs can potentially improve the selection of initial conditions for adaptive sampling. In recent years, there have been attempts in the recent years to extract CVs using machine learning (ML) methods. Unsupervised learning models for CV identification, which are trained on unlabeled data points (i.e., sampled configurations without additional information), have shown promise. Currently, the most widely used nonlinear ML dimensionality reduction method for CV discovery is autoassociative neural networks, also known as autoen-coders (AEs).^1^ Notable and effective examples include time-lagged independent component analysis (tICA), the variational approach for Markov processes (VAMP),^2^ VAMPNets,^3^ and time-lagged variational autoencoders (TVAE).^4^ These unsupervised models aim to recover patterns and intrinsic properties from the kinetic information of a set of parallel trajectories. Transformer architecture has been proven useful in encoding proteins into latent space to explore their conformational ensembles of protein through MD simulations.^5^ In this study, we proposed a Dynamic-Mixed transformer-based framework (DMFormer) to accelerate exploration of the conformational ensembles of allosteric protein Gαo. Gα is a subunit of heterotrimeric guanine-nucleotide-binding regulatory proteins (G proteins), which are crucial for signal ransduction.6 In its basal state, Gα is bound to guanosine diphosphate (GDP) and interacts with Gβγ subunits. The exchange of GDP for guanosine triphosphate (GTP) triggers the allosteric activation of Gα, enabling it to interact with and regulate downstream effector proteins.^7^ Gαo and Gαi are both classified into Gαi family based on sequence homology.^8^ However, the performance of Gαo remains unclear due to the lack of crystal structure. In this work, we aim to apply our DMFormer to analyze the allosteric pattern of Gαo.

First, the crystal structure of the Gαo complex was subjected to 500 ns MD production simulations. This process was repeated to obtain five trajectories. Then, the trajectory data were split into 11,250 ensembles (50 frames per ensemble after 50 ns) to serve as the training set for one system. We used the Transformer-based generative neural network to learn the conformational dynamic within the training set. This model was then used to generate a convincing interpretability of the allosteric mechanism.

## METHODS

### Molecule Dynamics Simulation

The initial structure of active and inactive Gα_o_ (Uniprot ID: P16378, residues 25-354) were obtained using AlphaFold2.^9^ These predictions were made for isolated G*α*_*o*_ and complex of Gα_o_ and G-protein Regulatory Motif (GoLoco), respectively (see FIG 1). The initial binding sites of Magnesium ions and GTP were identified by aligning Gα_o_ with the crystal structure of Gα_i1_ complex (PDB ID: 1GIA). Similarly, the binding sites of GDP were identified using the crystal structure (PDB ID: 1GDD). The system was neutralized by adding Na^+^ and Cl^−^ ions, and solvated in a 9 nm periodic bounding box of TIP3P water molecules.^10^ The topology files of the system were prepared using the CHARMM36m force field.^11^

**Figure 1.**
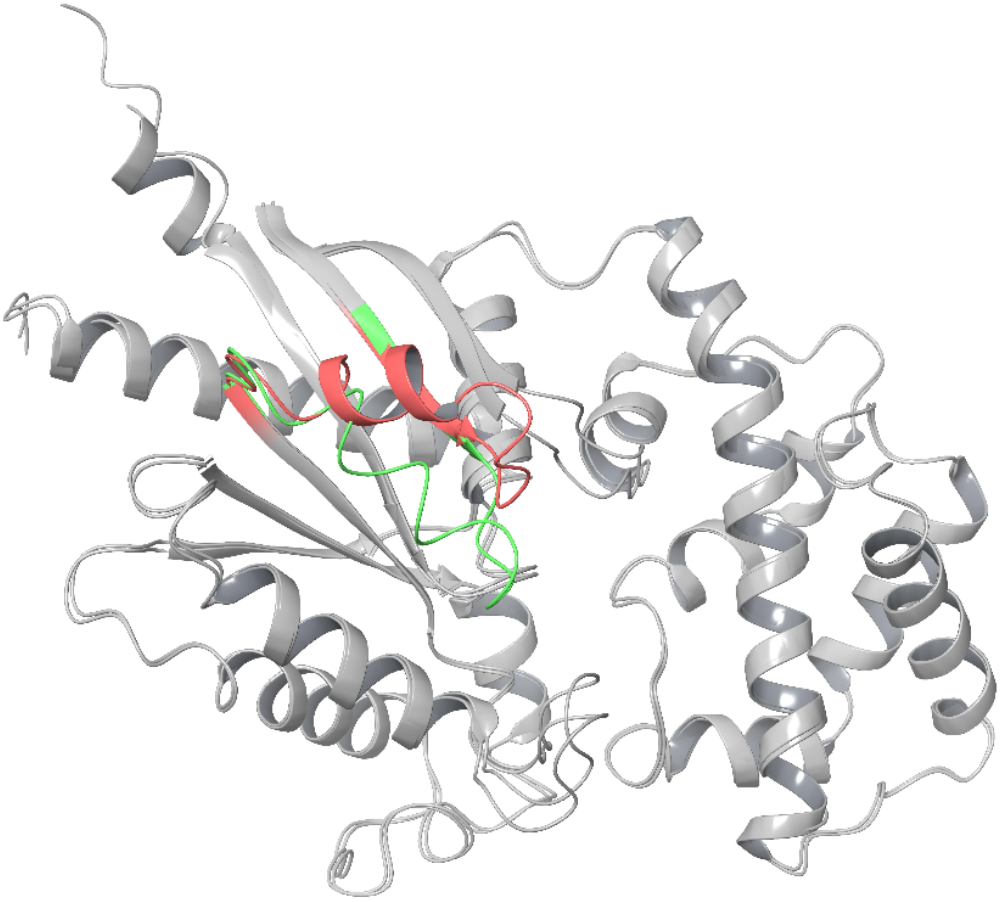
The Initial structure of active and inactive Gαo. The Switch II region of the active structure is colored green, while that of the inactive structure is colored red.

**Figure 2.**
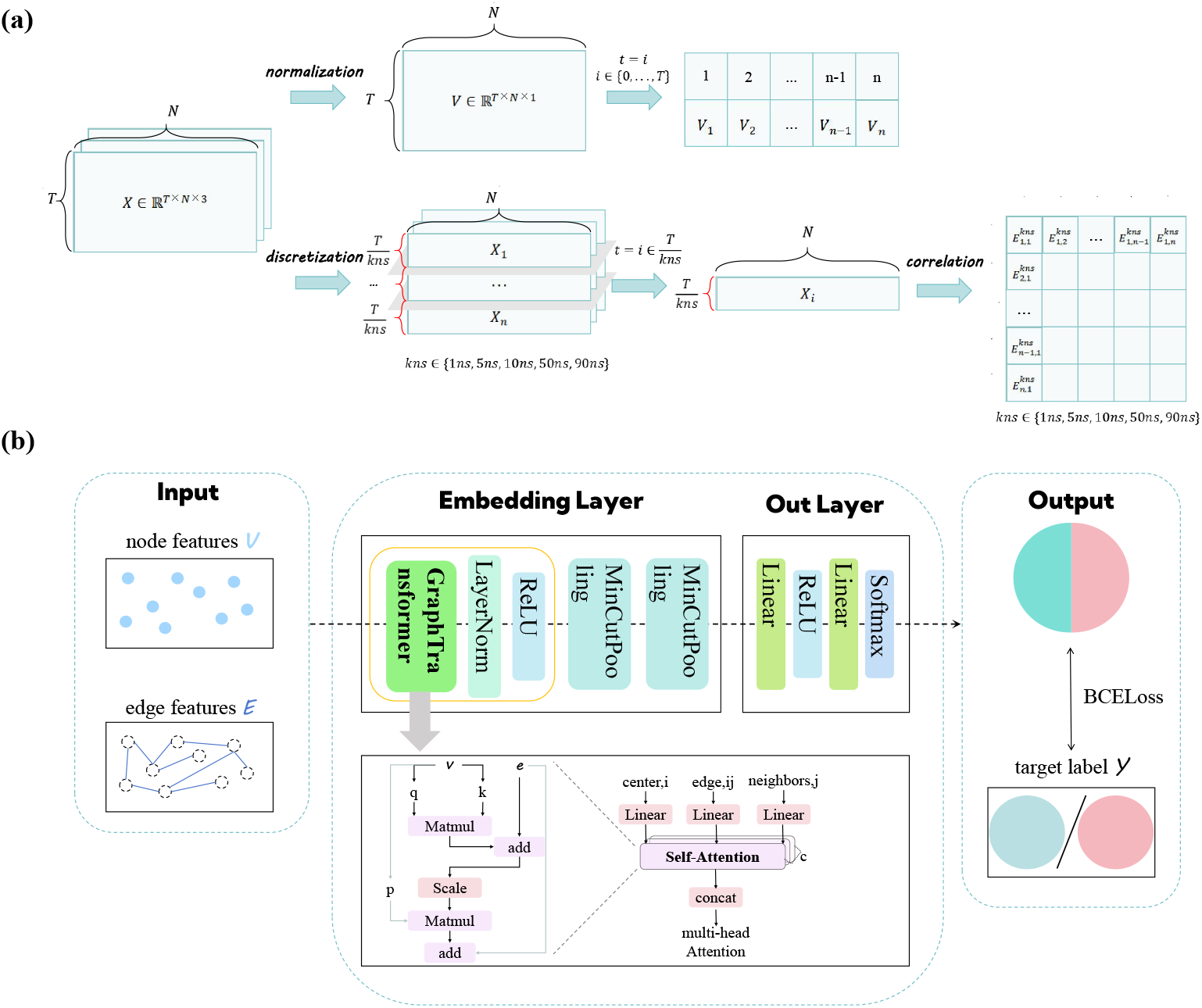
Pipeline for Model Training. (a) Workflow for processing and feature extraction from raw data. (b) Architecture of the model backbone. Interpreting Model Predictions Using Integrated Gradients

The simulated system was first energy minimized by performing a 50,000-step typical ensemble (NVT) simulation with the Berendsen integrator, and then further equilibrated by performing a 50,000-step NPT simulation at 310 K. Particle Mesh Ewald (PME) algorithm12 was applied for the calculation of long-range electrostatic interactions, with a cutoff of 1.2 nm for direct interactions. The SHAKE algorithm13 was set to constrain the lengths of all bonds involving hydrogen atoms. The simulation time step was set to 20 ps, followed by five parallel 500 ns MD productions were performed. All simulations were conducted using Gromacs software package.^14^

### Transformer-based Graph Neural Network

Recent experiments have demonstrated the applicability of graph-based neural networks in encoding proteins by utilizing graphical representation.^15–17^ This approach provides a natural spatial representation of molecular structures and is more interpretable than 2D volumetric representations. In this study, we proposed a supervised Transformer-based graph neural network backbone to depict protein ensembles.

### Graph representation of protein snapshot

In this work, we constructed a molecular graph for each protein snapshot, referred to as the amino-acids/residues motion network, to capture the dynamic information of the trajectories. Our method for extracting dynamic information from trajectories involves: (i) Choosing an appropriate vector representation for residue motions. (ii) Choosing an appropriate vector representation for the residue-residue cooperative motion patterns. Since protein motion patterns differ between slow and fast motions, we included residue-residue cooperative motion patterns in five timescales (1 ns, 5 ns, 10 ns, 50 ns, 90 ns).

First, the MD trajectories of the system were preprocessed to obtain the sequence length and displacement relative to the reference structure. Cβ atoms (Cαs for Glycine) were selected to represent the motion of the residues. Consider a trajectory *X* ∈ℝ^*T*×*N*×3^, where *T* denotes the total number of snapshots, taken at intervals of 20 over a duration of 450 ns. Here, *N* represents the number of residues, and the final dimension of 3 corresponds to the *x, y*, and *z* coordinates. After subtracting the mean value and normalizing this matrix *X* (as shown in Equation 1), we obtained the resulting matrix *V* ∈ℝ^*T* × *N* ×1^.

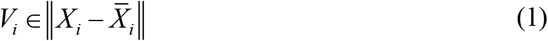

Second, five independent 450 ns-long MD trajectories (equilibrated after 50 ns) of each Gαo-ligand complex system were discretized into bins with duration of 1 ns, 5 ns, 10 ns, 50 ns, and 90 ns. The dynamic cross-correlation matrix *E* ∈ ℝ^*N*×*N*^ was calculated using the normalized covariance for each bin, and this matrix was shared by all snapshots within the same bin:

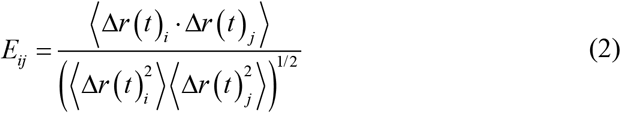

where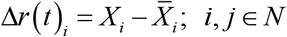 represents two residues. In this way, we assigned five dynamic cross-correlation matrices in different timescales to each snapshot.

As input for graph neural networks, we constructed a digraph *G* (*v, e*) for each snapshot in a system, where *v* is a set of nodes and *e* is a set of edges between every pair of distinct nodes. The nodes are described by the feature matrix *V* ∈ℝ ^*N* ×1^ and serial numbers ∈ℝ^*N*×1^for positional identification, while the edges are described by the feature matrix 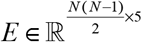. The adjacency matrix *A* = [*a*_*ij*_] ∈ ℝ^*N*×*N*^ is used to describe the graph *G*, and the diagonal degree matrix is denoted by *D ∈ diag* (*d*_1_, *d*_2_,…*d*_*n*_), where 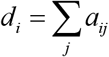 is the degree of node *i*. We sampled data from the entire snapshots at intervals of 50 to form the dataset for a system.

### Model Architecture

In this section, we introduce our DMFormer framework, designed to learn features from data represented as graphs. The DMFormer model enhanced the analysis of MD simulation trajectories by leveraging advanced deep learning techniques. The primary objective of DMFormer is to accurately capture and represent the intricate spatiotemporal patterns within these simulations, facilitating more insightful and precise scientific analysis.

The proposed model features an encoding block that comprises a TransformerConv^18^ layer followed by two pooling layers, and an output layer consisting of a linear transformation layer. The TransformerConv layer is designed to process both node features and edge features within the graph, transforming these into node classifications. This transformation effectively captures the intricate relationships and dependencies between the nodes and edges.

Following the TransformerConv layer, each of the two pooling layers sequentially reduces the number of nodes by half, aiding in distilling the most salient features and reducing computational complexity. These pooling operations condense and distill critical information from the high-dimensional MD data. After the two pooling layers, an averaging operation is applied to the nodes, reducing the number of nodes to one. This operation aggregates the node features, providing a comprehensive representation of the entire graph.

Finally, the output layer employs a linear transformation to convert the aggregated node features into a single numerical value. This value serves as the final prediction, enabling the model to perform graph classification with high accuracy. Through this architecture, the model effectively integrates advanced feature extraction and dimensionality reduction techniques to enhance its predictive capabilities on graph-structured data.

### Graph TransformerConv

The TransformerConv layer plays a crucial role in capturing long-range dependencies and interactions within the molecular simulation data. By leveraging self-attention mechanisms, it can discern and highlight the most relevant features and patterns throughout the dataset. This capability is vital for understanding the intricate behaviors and properties of molecules over time.

Graph Transformer Conv (GTC) transforms and propagates node features *X* and edge features *E* across the graph by several layers, including linear layers and nonlinear activation to build the approximation of the mapping: *X* → *Y*, where *Y* is the node class matrix.

The feature propagation scheme of GTC in layer *l* is: Given source features 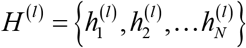, we calculate multi-head attention for each edge from *j* to *i* as following:

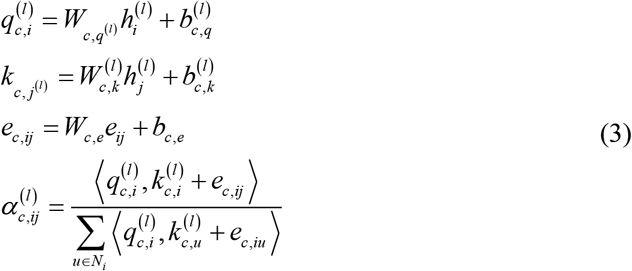

where 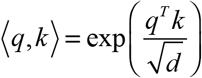 is exponential scale dot-product function and *d* is the hidden size of each head, and *N*_*i*_ are the neighbor nodes of *i*. For the *c*-th head attention, we firstly transform the node features of node *i* and *j* into query vector 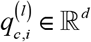 and 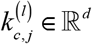 respectively using different trainable parameters 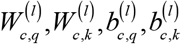. The provided edge features *e*_*ij*_ will be encoded and added into key vector as additional information. When *l* = 0, *H*^(0)^ is assigned to the node features *V*. After getting the graph multi-head attention, we make a message aggregation from the distant *j* to the source *i*:

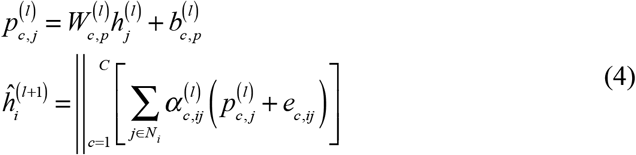

where the is the concatenation operation for *C* head attention. The distant feature *v*_j_ is transformed to 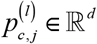 for weighted sum.

In addition, we add a gated residual connection following the layer as shown in Equation 5 to prevent our model from over smoothing.

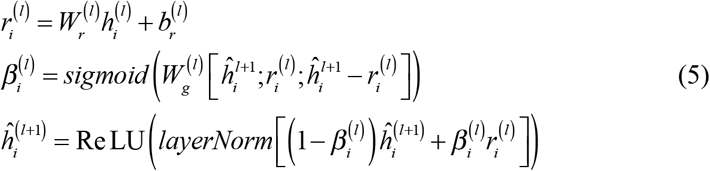

### Model Training and Hyperparameters

Our model is initialized with specific parameters, including the number of input channels, hidden channels, average nodes, output channels, and the number of heads in the Transformer convolutional layer. These parameters are set to 2, 16, 330, 1, and 1, respectively.

The training set included datasets of Gα_o_(active)-GTP-Mg^2+^ and Gα_o_(inactive)-GDP systems, assigned as positive and negative labels, respectively. To study the process of conformational changes, we aimed to observe the performance of these two systems in the training set after ligand exchange. Therefore, the test set included datasets of Gαo(inactive)-GTP-Mg^2+^ and Gαo(active)-GDP systems, which were expected to have opposite labels, negative and positive, respectively. For the Q205L mutant, we replicated the conditions used for the wild type systems to form test sets.

The training consisted of 10 epochs, and bath size was 64. To optimize the model, we used the Adam optimizer (learning rate = 0.001), and the loss function is Binary Cross Entropy Loss (BCELoss):

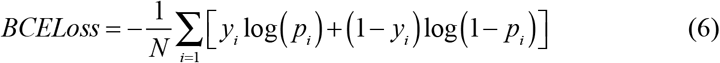

where *p* represents the propensity of a conformation to become activated. A value of 1 indicates the active state, while a value of 0 indicates the inactive state. The evaluation metric used was the AUC score.

To enhance the interpretability of our model, we employed the Integrated Gradients method,^21^ a widely recognized approach for attributing the predictions of deep learning models to their input features. Integrated Gradients (IG) address the need for understanding how individual features contribute to the model’s output, providing a more transparent and interpretable prediction process.

The core idea of IG is to quantify the contribution of each input feature by integrating the gradients of the model’s output with respect to the input along a path from a baseline input to the actual input. First, a baseline input is selected (usually zero), which represents the absence of features with the same dimension of the input features. Next, a series of interpolated inputs are generated between the baseline and the actual input. This interpolation is performed in small steps along the path from the baseline to the input, capturing intermediate states. For each interpolated input, the gradient of the model’s output with respect to the input is computed. This gradient indicates how changes in the input affect the output at that specific point.

The gradients are then integrated (summed) along the path of interpolated inputs. For an input × and a baseline *x*’, the integrated gradient for the *i*-th feature is computed as:

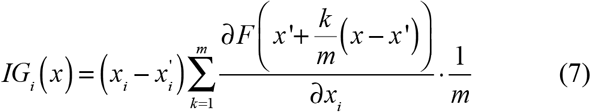

where *F* represents the model’s output function, *m* is the number of steps in the interpolation, and *k* is the step index. Mathematically, this involves accumulating the gradients at each step and multiplying by the step size, effectively capturing the total change in the output due to the change in each input feature from the baseline to the actual input. The integrated gradients provide attribution scores for each input feature, reflecting their contribution to the model’s prediction. These scores can be visualized to highlight the most influential features. For ease of comparison, we normalized the IGs of different systems using the global extrema to obtain weighted IGs.

## Result and Discussion

We analyzed the prediction results of the DMFormer on both the training set and test set. To visually present the prediction outcomes, we plotted the predicted values as line charts over time. This approach not only reveals the temporal trends of protein conformational changes but also allows us to observe the dynamic transitions between active and inactive states. Through an in-depth analysis of each trajectory, we further quantified the proportions of active and inactive conformations. This detailed examination provided insights into the frequency and conditions under which each state occurs, contributing to our understanding of the dynamic behavior of the protein systems under study.

### DMFormer Performance on Training Set

**T**he prediction results on the training set aligned well with the target prediction outcomes, achieving an AUC of 0.75. This indicates that the model has been well-trained on the training set, demonstrating its effectiveness in capturing the desired predictive patterns. In the stable inactive system Gαo(inactive)-GDP, all of the five trajectories had inactive conformations exceeding 50%, while one trajectory consistently remained in the origin conformation (see TABLE 1), resulting in an average inactive proportion of 68% (see FIG.3a). Similarly, in the stable active system Gαo(active)-GTP-Mg^2+^, the sections in all five parallel trajectories were close to active conformation, with the average activation proportion reaching 90% (see FIG.3b).

**Table 1:** Percentages of active state in Gα_o_ systems with different nucleotide-bound.

| Traj. | Gαo(inactive)GDP | Gαo(active)GTP | Gαo(inactive)GTP | Gαo(active)GDP |
| --- | --- | --- | --- | --- |
| 1 | 49% | 72% | 83% | 86% |
| 2 | 31% | 88% | 91% | 82% |
| 3 | 0% | 99% | 81% | 91% |
| 4 | 43% | 97% | 80% | 59% |
| 5 | 36% | 92% | 87% | 32% |
| Average | 32% | 90% | 84% | 70% |

This dynamic behavior is likely a prerequisite for the protein’s response to external signals, allowing it to rapidly transition to an active state upon receiving the GTP signal. This suggests a conformational selection binding mechanism. Our experimental results support the conformational selection model, indicating that the Gαo protein can retain the potential to stay in an active state even when bound to GDP. This “pre-activated” state may provide a biological basis for a quick response to external signals.

### DMFormer Performance on Test Set

We exchanged the ligands of two stable systems and anticipate that the conformation of Gαo in these two unstable systems would transition from their original state to the other state (see Table 1). In the Gαo(inactive)-GTP-Mg2+ system, after 50 ns of equilibration, the conformation of Gαo remained predominantly in the active state (see FIG.3c). Across the five parallel trajectories, the proportion of active conformations was greater than 80%.

**Figure 3.**
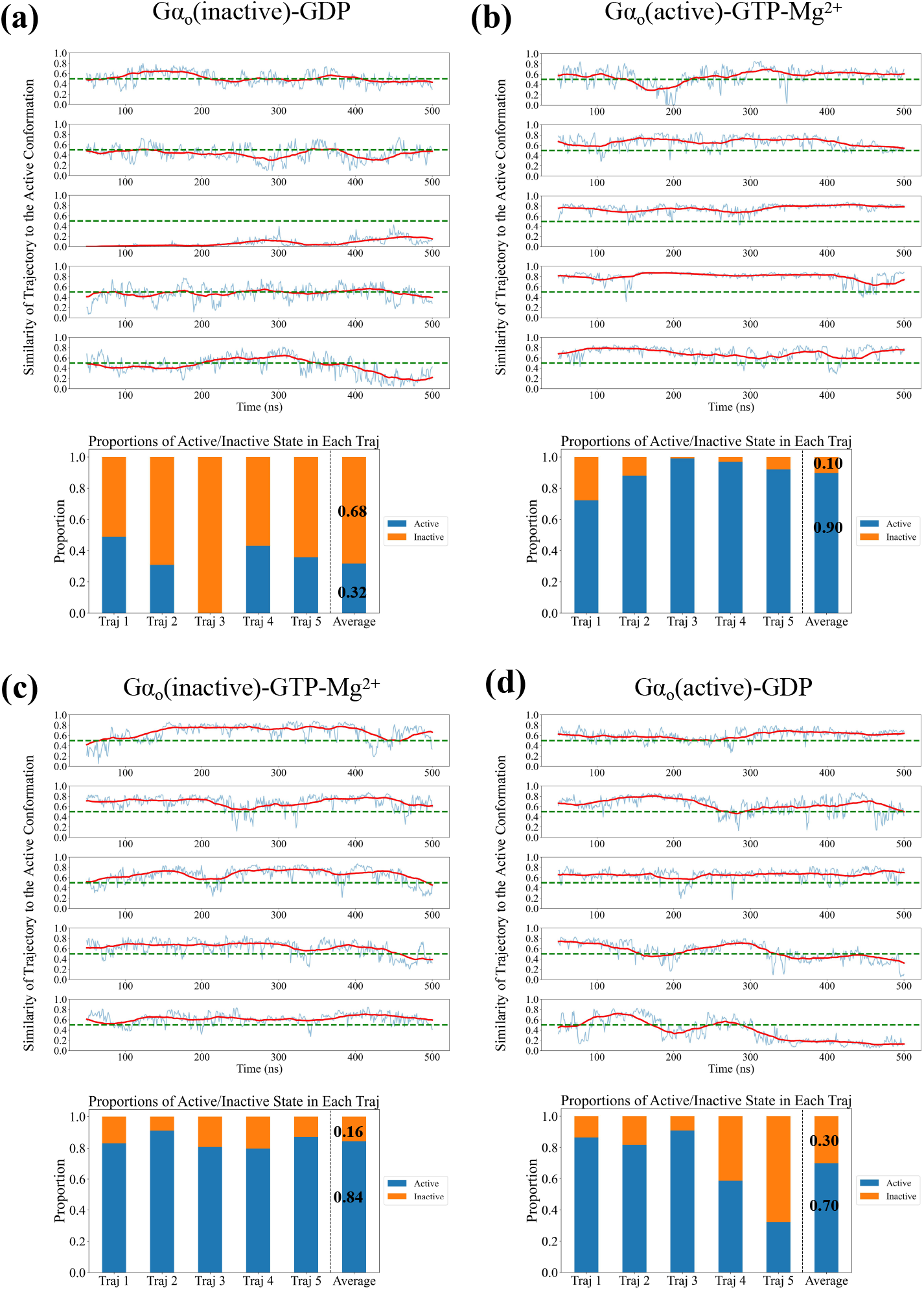
Predicted state of each section in trajectories along the simulation and proportions of active and inactive states in each trajectory for (a) Gαo(inactive)-GDP, (b) Gαo(active)-GTP-Mg^2+^, (c) Gαo(inactive)-GTP-Mg^2+^, (d) Gαo(active)-GDP.

Overall, 84% of the conformations were in the active state, which is a significant increase compared to the Gαo(inactive) system bound to GDP. This aligns with our hypothesis that the conformation of inactive Gαo is easily regulated by GTP, transitioning into the active conformation.

In the Gαo(active)-GDP system, we observed that after ligand exchange, the conformation of Gαo gradually transitions from the initial active state to an inactive state over the simulation time (see FIG.3d). In three out of the five parallel trajectories, the predicted activation probability shows a significant downward trend over time, suggesting the process of active Gαo hydrolyzing GTP to GDP and subsequently reverting to the inactive state. Overall, 30% of the conformations reverted to the inactive state post-GTP hydrolysis, but the transition process was relatively slow. By analyzing the time-dependent conformational prediction probability curves, we found that the system’s conformation state was slowly changing during the 500 ns simulation time in the last two trajectories, without reaching a stable state, which can also be observed from the RMSD curves (see FIG.4). Therefore, we believe that with a longer simulation time, the proportion of inactive conformations will further increase. However, even in a shorter simulation, we could already capture the trend of the active conformation transitioning to an inactive conformation.

**Figure 4.**
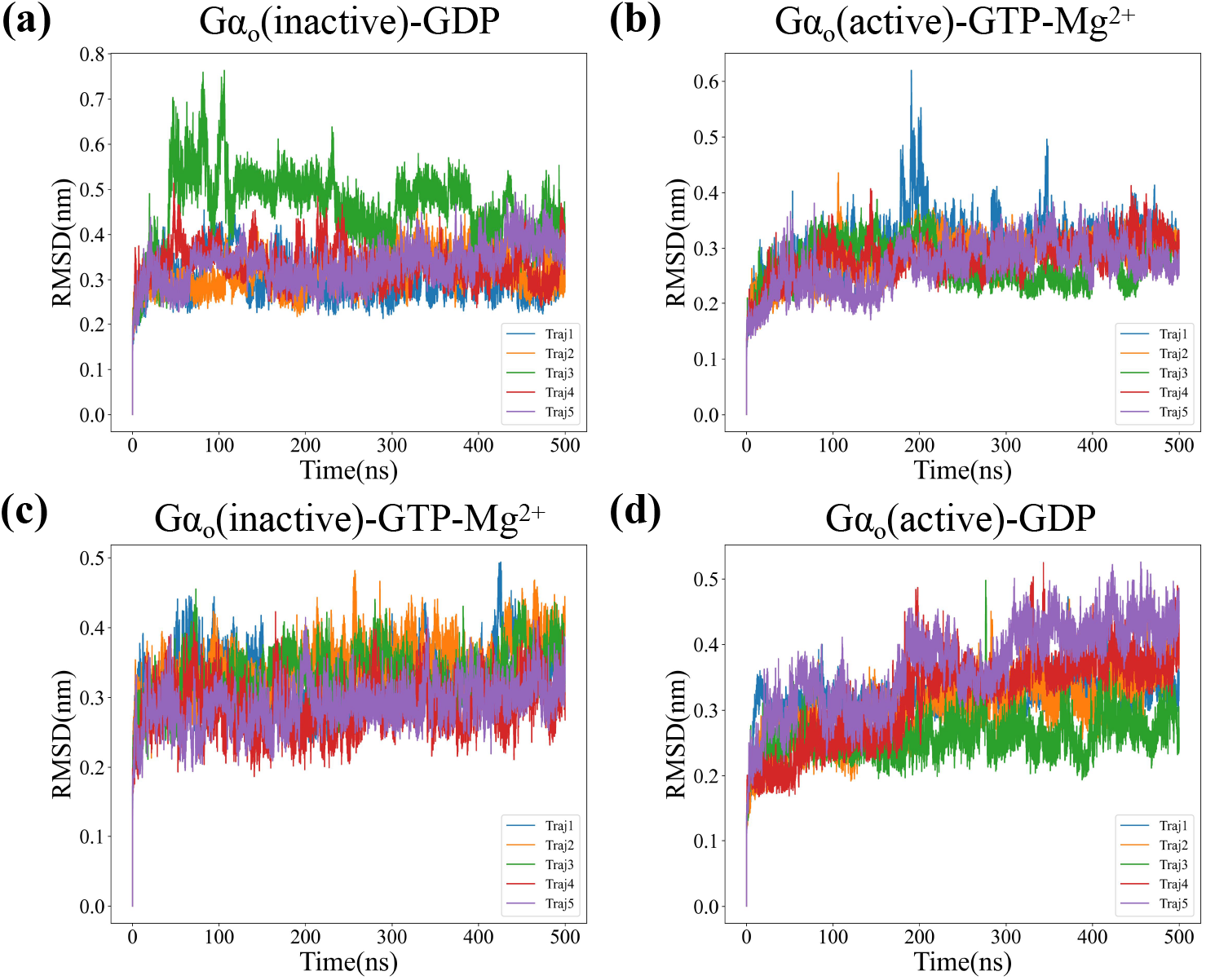
Root-Mean-Square Deviation (RMSD) Compared to Initial Structure During the Simulation for Wild Type Systems.

Next, we aimed to validate the predictive performance of this model on mutants (see FIG.5). We selected mutant Gαo-Q205L, which remains constitutively active. In the Gαo-Q205L system, the inactive state is missing. However, to facilitate a more straightforward comparison, we artificially constructed the missing states in this system. For the Gαo-Q205L system, we found that 86% of the Q205L mutant’s active conformations remained in the active state after binding with GTP (see TABLE 2).

**Figure 5.**
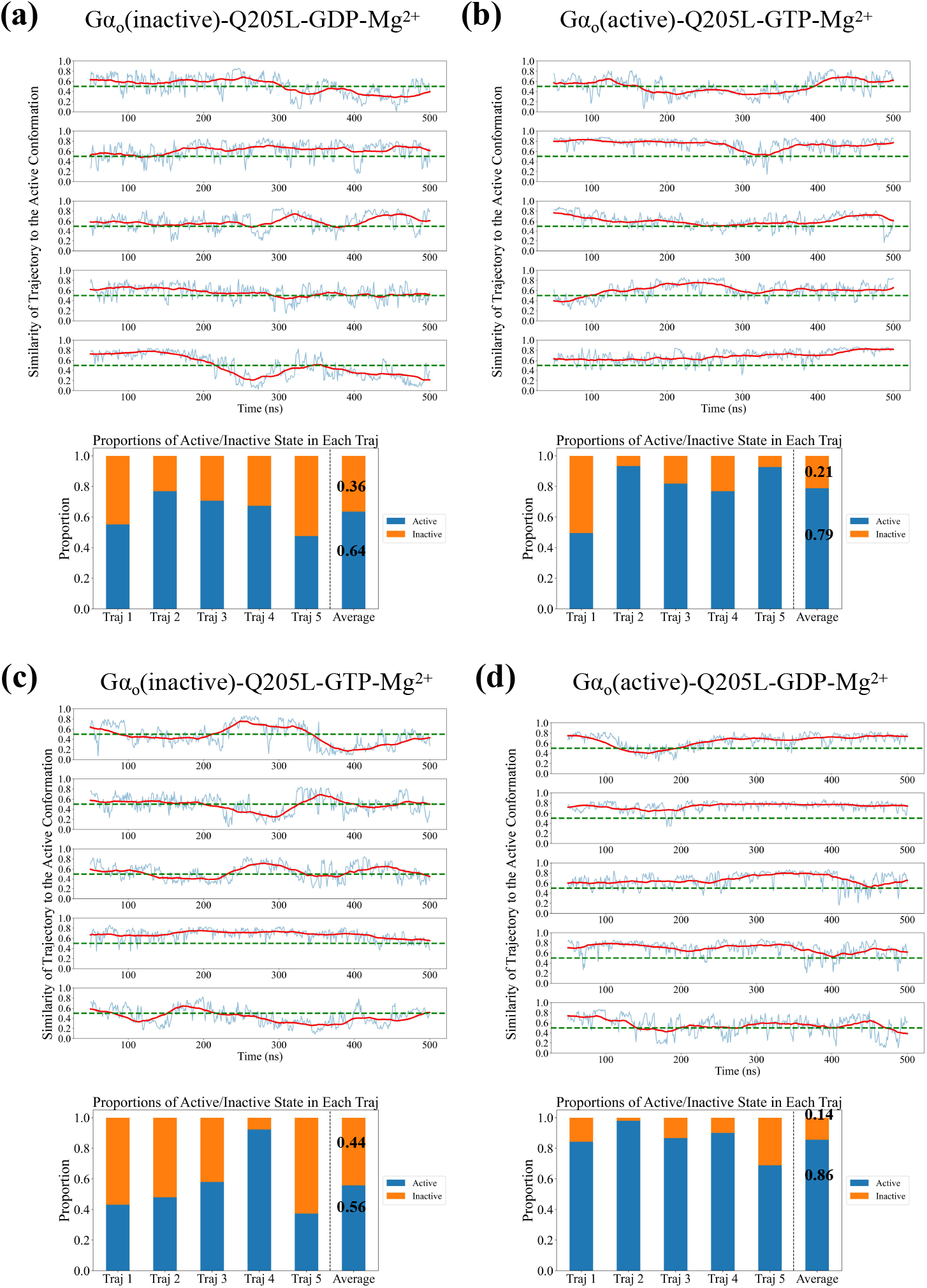
Predicted State of Each Section in Trajectories Along the Simulation and Proportions of Active and Inactive States in Each Trajectory for (a) Gαo(inactive)-Q205LGDP-Mg^2+^ (b) Gαo(active)-Q205L-GTP-Mg^2+^ (c) Gαo(inactive)-Q205L-GTP-Mg^2+^ (d) Gαo(active)-Q205L-GDP-Mg^2+^.

**Table 2:** Percentages of active state in Q205L mutant systems with different nucleotide-bound.

| Traj. | Q205L(inactive)<br>GDP | Q205L(active)<br>GTP | Q205L(inactive)<br>GTP | Q205L(active)<br>GDP |
| --- | --- | --- | --- | --- |
| 1 | 55% | 84% | 43% | 50% |
| 2 | 77% | 98% | 48% | 93% |
| 3 | 71% | 87% | 58% | 82% |
| 4 | 67% | 90% | 92% | 77% |
| 5 | 48% | 69% | 37% | 93% |
| Ave. | 64% | 86% | 56% | 79% |

Additionally, in the artificially constructed inactive conformations of the mutant, 64% transitioned to the active state upon binding with GDP (see TABLE 2). This indicates that, regardless of the initial state, the Q205L mutant favors the active state. After ligand exchange, we observed that 79% of the Q205L mutant’s active conformations remained in the original state even after binding with GDP (see TABLE 2). This demonstrates that the Q205L mutant’s active conformation is not affected by ligand binding and remains constitutively active. These findings are consistent with experimental results, showcasing the model’s ability to generalize and explain the conformational changes in mutants.

### Interpretability of DMFormer

To elucidate the dynamic mechanisms underlying Gαo protein conformational changes, we analyzed the influence of input features (including node and edge features) on the output results using the Integrated Gradients method. We normalized the Integrated Gradients of both node and edge features based on global extrema to facilitate comparison. A higher positive value of the weighted Integrated Gradient for a feature suggests that an increase in this feature’s value enhances the probability of predicting the conformation as an active state. Conversely, a lower negative value of the weighted Integrated Gradient indicates that an increase in this feature’s value raises the likelihood of predicting the conformation as an inactive state. In our dataset, node features represent the displacement distance of a residue relative to the reference structure, while edge features denote the correlation of displacement between residue pairs. Thus, a larger node feature value indicates greater flexibility of that residue, and a larger edge feature value reflects stronger cooperative motion between residue pairs.

### IGs of Node Features

First, we analyzed the impact of node features on conformation state prediction in the training set. We found that the weighted Integrated Gradients of all node features were negative (see FIG.6a), indicating that all residues of the Gαo protein exhibit lower flexibility and higher stability in the active state. We observed several prominent peaks in the residue weighted Integrated Gradient negative value curve, corresponding to residues 25-32, 32-72, 105-122, 178-190, 199-221, 232-243, and 257-263 of the Gαo protein. By aligning the structure of the Gαo protein with the crystal structure of the Gαi1 protein (see FIG.6b,c), we identified that these amino acid residue segments belong to the αN domain, Switch I domain, Switch II domain, Switch III domain, and the loop segment between α9 and β5 (referred to as the α9 − β5 loop), respectively.

**Figure 6.**
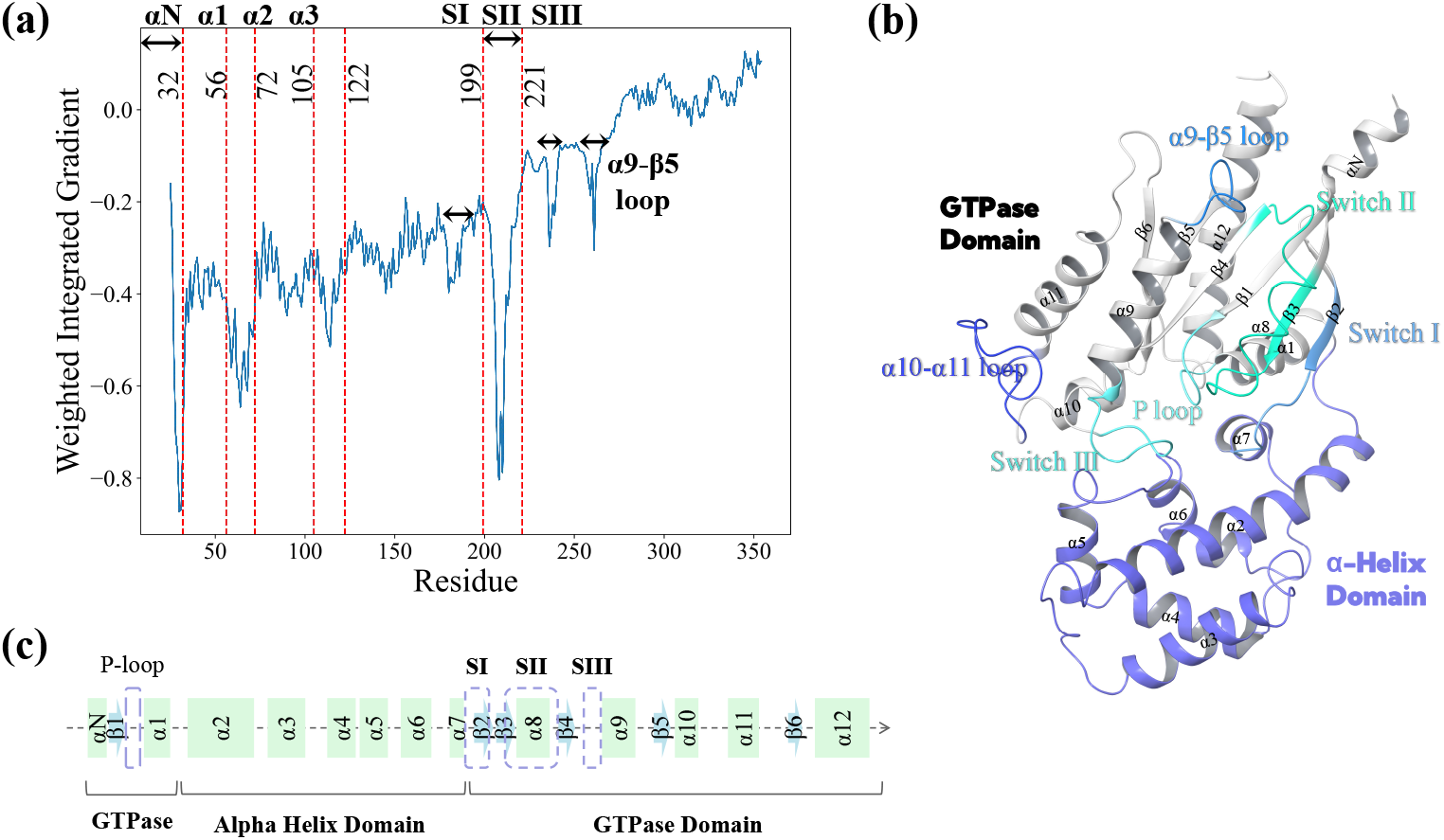
Interpretability based on node features. (a) Weighted Integrated Gradient of Node Features from the Training Set. Gαo Structure Labels in (b) 3D Space and (c) Sequence.

Among them, the peak value of the weighted Integrated Gradient corresponding to the Switch II domain is the highest. This indicates that the residue movements in this segment differ significantly between the active and inactive states, making it a key marker for the model to distinguish between these two states. In the active state, the Switch II domain exhibits smaller movements and is more tightly integrated with the overall structure. Switch I, Switch III, and the α9-β5 loop segments also correspond to higher weighted Integrated Gradient values, indicating that these segments are also stable in the active state. Based on previous analysis of the Gαi1 protein,^23,24^ earlier studies have found that these segments play a crucial role in the stable binding of Gα proteins with GTP.

Therefore, we believe that the interpretability analysis provided by the model based on node features has practical biochemical significance. It can guide us in identifying key segments that play important roles in the conformational changes of the protein.

### IGs of Edge Features

Next, we analyzed the influence of edge features on the prediction of the active state by calculating their integrated gradients. Given that our edge features have five channels, each representing a different time scale, we first analyzed the average integrated gradients of edge features across these five channels. This approach helps us capture the most significant patterns of cooperative motion between residues.

When the weighted integrated gradient is positive or negative, it indicates that the feature contributes to the prediction of the active or inactive conformation, respectively. Therefore, we divided the weighted integrated gradients of edge features into positive (red, see fig.7a) and negative (blue, see fig.7b) parts. The deeper the color, the larger the absolute value of the gradient, indicating a greater influence on the conformation prediction structure. We found that the correlation coefficients of movements within residues 25-32 (αN) provide a very significant positive contribution, while the correlation coefficients of movements between αN and residues 199-221 (Switch II) provide a very significant negative contribution. This indicates that in the active conformation, the internal movement correlation of αN is higher compared to the inactive conformation, potentially representing greater structural stability. Conversely, in the inactive conformation, there was a strong correlated movement between αN and Switch II, which may suggest an indirect interaction between these regions.

**Figure 7.**
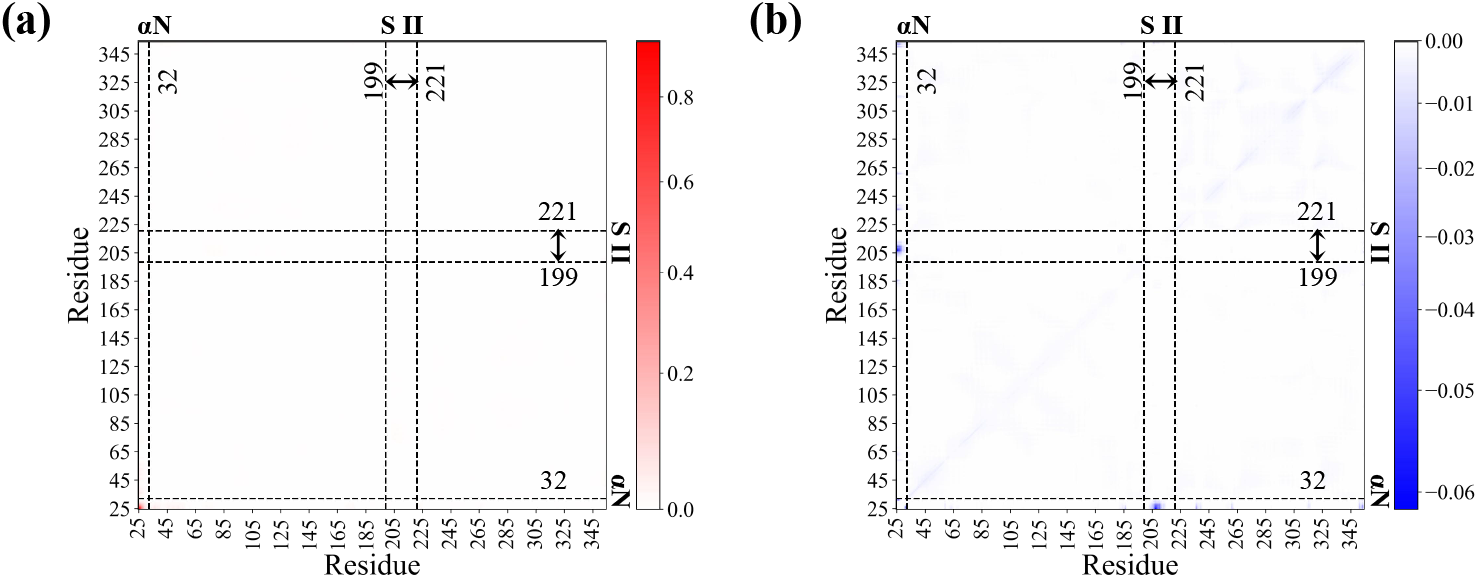
Interpretability Based on Edge Features. Positive Integrated Gradient in Red (a) and Negative Integrated Gradient in Blue (b).

We also observed the correlated movement patterns between segments other than αN (see FIG.8a,b). Based on the structure, we divided the residue sequences into the GTPase domain and the α-Helix domain. The GTPase domain hydrolyzes GTP to GDP, while the α-Helix domain plays an important regulatory role. We found that in the active conformation, there is a close correlated movement between the GTPase domain and the α-Helix domain (see FIG.8a). In the active state, several segments within the α-Helix domain exhibit correlated movement with the entire GTPase domain. Among these, the correlated movement pattern between the Switch II segment and the GTPase domain is the most pronounced and significant. In contrast, in the inactive conformation, the two domains act as independent entities, exhibiting more internal correlated movements within each domain (see FIG.8b). Notably, the internal correlated movement patterns of the α9-12 and β5-6 segments make significant contributions.

Through weighted integrated gradient analysis over different time scales, we identified distinct movement patterns at various speeds (see FIG.8c). Our analysis revealed that the integrated gradient of the correlated movement coefficient between the GTPase domain and the α-Helix domain is highest on the 50 ns time scale. This suggests that the correlated movement between these domains primarily occured as a slow movement on this timescale. Similarly, the correlated movement within the α-Helix domain also predominantly occurred on the 50 ns time scale. In contrast, the internal correlated movement within the GTPase domain was faster, predominantly occurring on the 5 ns time scale. We speculate that the GTPase domain is structured to respond rapidly to changes, whereas the α-Helix domain, due to its structural stability and longer sequence, requires more time to respond to signals. During the process of the α-Helix domain receiving external changes, it transmits signals to the GTPase domain through slow correlated movements, thereby regulating the activity of the Gα protein.

**Figure 8.**
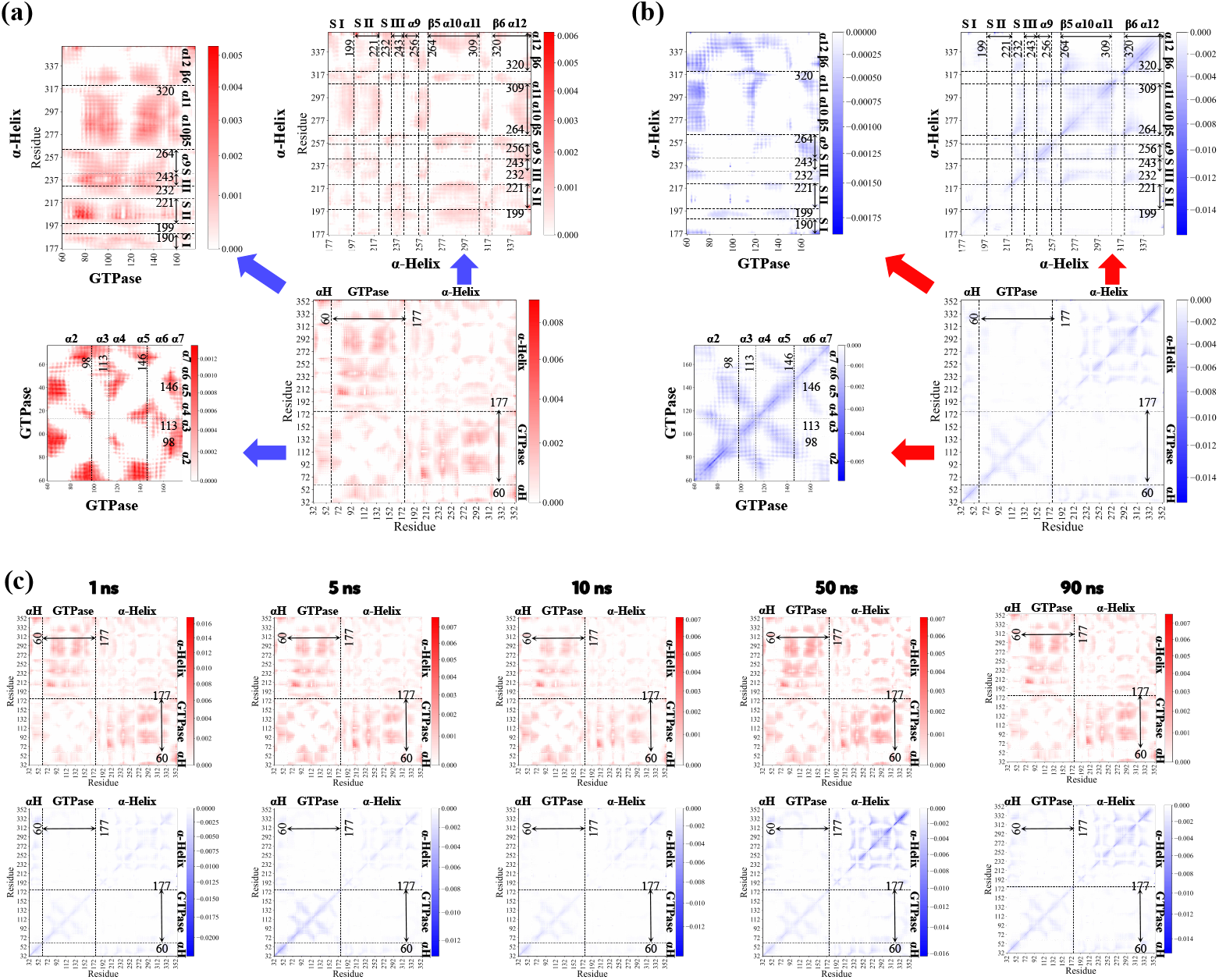
Interpretability based on edge features. Positive integrated gradient in red (a) and negative integrated gradient in blue (b). (c) Integrated gradients (IGs) of edge features at five different timescales, with positive integrated gradient in red and negative integrated gradient in blue.

### Explanation for Q205L Mutant

The analysis of node feature gradients provided a compelling explanation for the persistent activation of the Gαo Q205L mutant (see FIG.9a). The weighted integrated gradients derived from models based on the Gαo(inactive)-Q205L-GDP-Mg^2+^ and Gαo(active)-Q205L-GTPMg^2+^ datasets revealed a significant reduction in the absolute value of the integrated gradient for the Switch II segment in the Q205L mutant compared to the wild type systems. Moreover, there was a marked decrease in the absolute value of the weighted integrated gradients for both the αN segment and the residues 56-72 segment. This finding indicates that the reduced flexibility of the Switch II segment in the Q205L mutant underlies its persistent activation. The increased stability of Switch II prevents the mutant from reverting from the active conformation to the inactive state.

**Figure 9.**
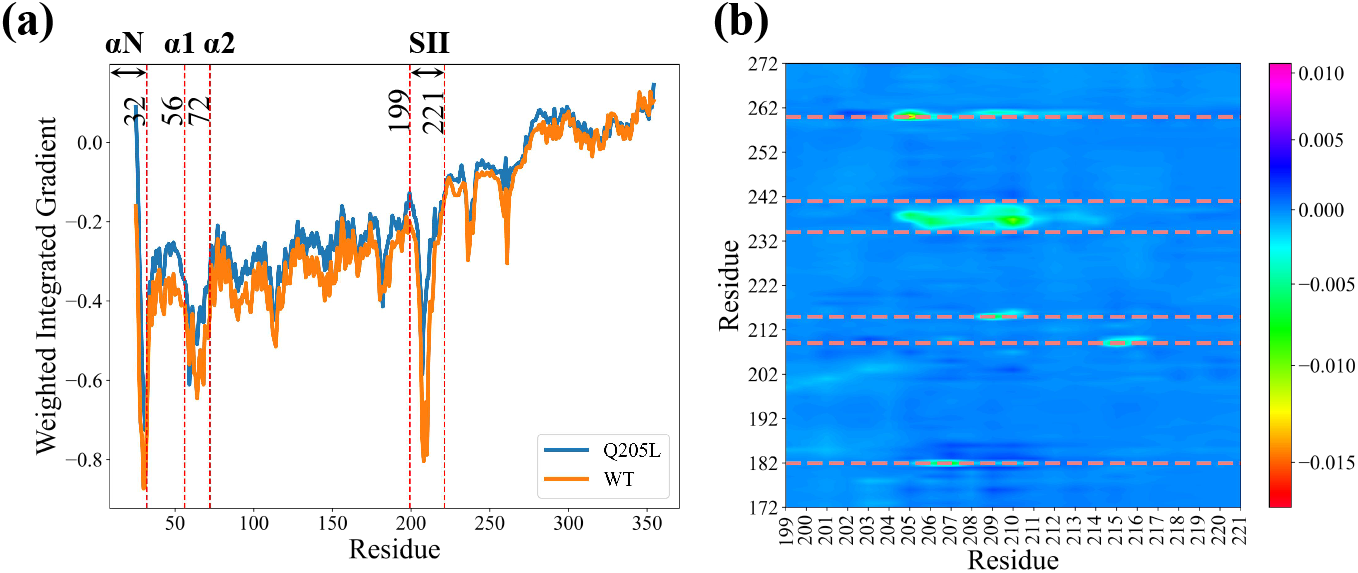
Interpretability based on node features (a) and Edge Features (b) of Q205L systems compared to wild type systems.

In the Q205L system, the weighted integrated gradients of edge features showed the most significant changes in the Switch II segment compared to the wild type (see FIG.9b). There was a notable decrease in the weighted integrated gradients of correlated movements between the Switch II segment and residues 182, 209, 215, 260, and residues 234-241 segment, presenting as negative contributions. In the wild type, the correlated movement patterns of this segment show a slight positive contribution to the weighted integrated gradient. This indicates that in the Q205L system, the movement patterns between these segments and the Switch II segment have altered, exhibiting an opposite trend compared to the wild type.

To better understand the impact of changes in edge features, we compared the differences between the wild-type system and the Q205L mutant system. We averaged the edge features of all samples in one system, as well as the edge features across five channels, to obtain the average distribution of edge features between residues within a system. We divided the entire protein sequence into eight bins based on the residue sequence, with each bin representing a node. The residues were grouped into the following independent nodes: 25-32, 33-60, 61-177, 178-198, 199-221, 222-243, 244-264, 265-354. These segments correspond to the domains ‘αN’, ‘α-Helix-1’, ‘GTPase’, ‘Switch I’, ‘Switch II’, ‘Switch III’, ‘α9-β5 loop’, ‘α-Helix-2’. The average value of edge features between nodes was used as the weight of the edges between nodes, constructing an undirected weighted graph. We displayed all edges in the graph with absolute weights greater than 0.1. The graph represents the movement correlation between internal domains of the Gαo protein, with positive correlations and negative correlations indicated by red and blue edges, respectively (see in FIG.10). The movement correlation graph of the system contains the communication pathways between different parts of the protein.

We found that the movement patterns of systems Gαo(active)-GTP-Mg^2+^ (see FIG.10b), Gαo(inactive)-GDP-Mg^2+^ (see FIG.10c), and Gαo(active)-GTP-Mg^2+^ (see FIG.10d) are essentially the same. This indicates that the information pathways in the Q205L system were consistent with those in the stable activation conformation of the wild type, further explaining the reason for the persistent activation of the Q205L mutant. In the Q205L mutant, even when the inactive conformation binds GDP, a significant portion of the conformations still transitioned to the active state. This also provides a reliable basis for the accuracy of our model.

**Figure 10.**
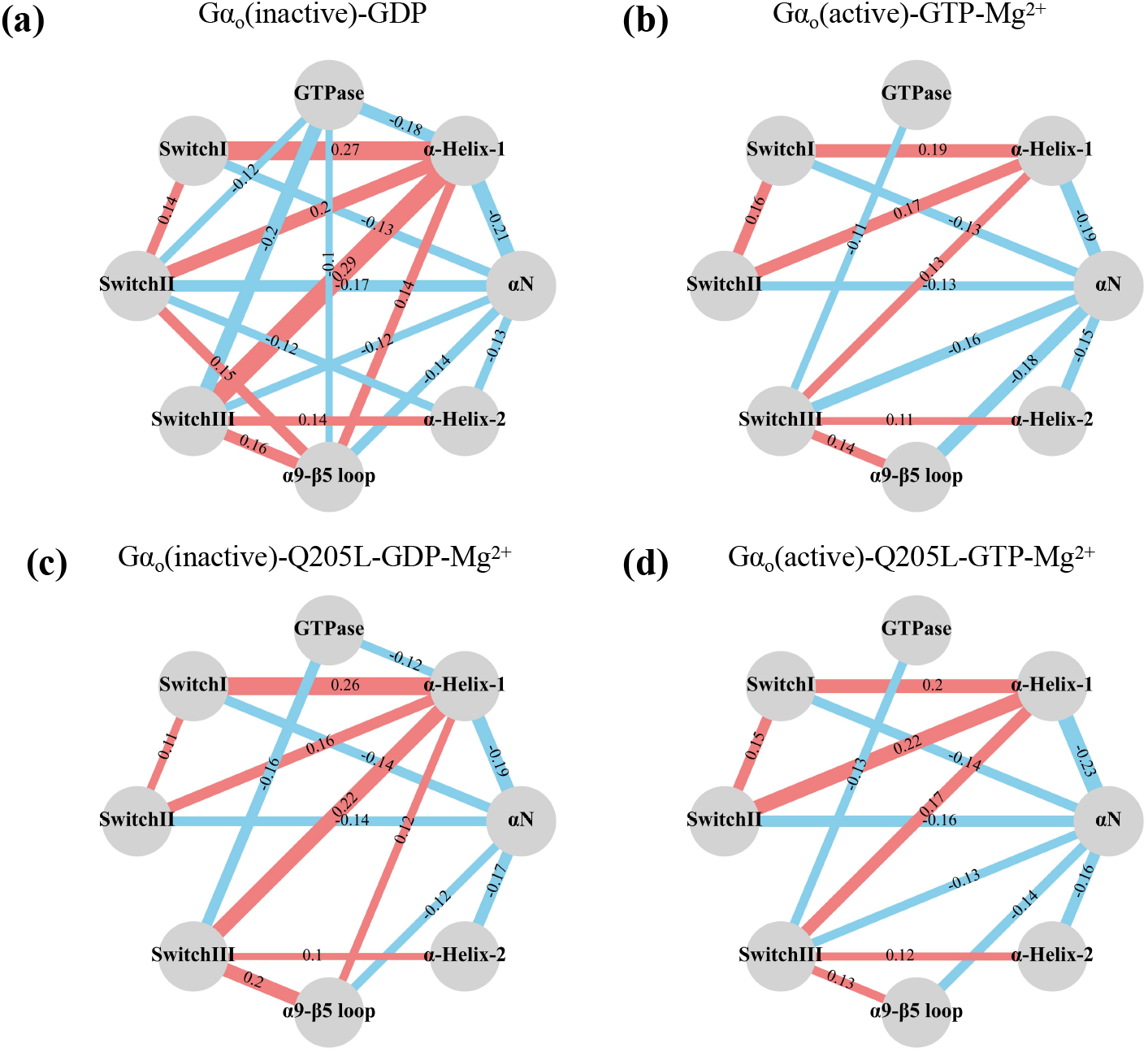
Undirected Weighted Graph Based on the Correlation Between Domains of (a) Gαo(inactive)-GDP, (b) Gαo(active)-GTP-Mg^2+^, (c) Gαo(inactive)-GTP-Mg^2+^, (d) Gαo(active)-GDP.

We found that in the Gαo(inactive)-GDP system, the values of the movement correlation coefficients between the Switch II and GTPase domain, Switch II and α9-β5 loop, and Switch II and α-Helix-2 were lower compared to the other three systems. Additionally, the values of the movement correlation coefficients between Switch II and α9-β5 loop were also higher in the Gαo(inactive)-GDP system. Further combining the weighted integrated gradient analysis of edge features, we found that the communication pathway between the GTPase domain and the α-Helix domain is crucial for determining whether the conformation is activated. Notably, the Switch II segment, which may act as a bridge facilitating communication between the Switch II and GTPase segments. In the Q205L mutant, the signaling capability of the Switch II segment is rigid, preventing the Gα protein from reverting to the inactive state. This impaired signaling likely explains the persistent activation observed in the Q205L mutant. The greater stability of Switch II and the higher correlation coefficients between the movements of Switch II and other segments suggest that Switch II is more stably bound to GTPase in the mutants.

## Conclusion

In this study, we developed and utilized a Transformer-based graph neural network framework, referred to as DMFormer, to investigate the conformational dynamics of the Gαo protein and its mutants. Our goal was to elucidate the mechanisms underlying protein activation and inactivation, focusing on the persistent activation of the Q205L mutant. The model’s predictive performance was validated on both wild-type and mutant systems, demonstrating robustness and generalizability.

The DMFormer model achieved an AUC of 0.75 on the training set, indicating its effectiveness in capturing the necessary patterns for distinguishing active and inactive states of Gαo. In stable inactive and active systems, the majority of trajectories maintained expected conformations, validating the model’s accuracy.

Analysis of ligand exchange experiments provided insights into conformational transitions. In the Gαo(inactive)-GTP-Mg2+ system, a significant proportion of conformations transitioned to the active state, highlighting the regulatory role of GTP. In the Gαo(active)-GDP system, a gradual transition to the inactive state was observed, indicating that GDP binding alone does not immediately revert the protein to its inactive state.

The interpretability of DMFormer was enhanced using Integrated Gradients. Node feature analysis identified key segments, such as the Switch II domain, as critical stabilizers in the active state. Edge feature analysis revealed communication pathways within the protein. Our study demonstrated that the GTPase and α-Helix domains exhibit different movement patterns depending on the protein’s state. In the active conformation, these domains showed strong correlated movements, particularly involving the Switch II segment. In contrast, the inactive state displayed more independent domain behavior, with pronounced internal correlations.

By averaging edge features across multiple channels and constructing an undirected weighted graph, we visualized the communication pathways within Gαo. The graphs highlighted both positive and negative correlations between various domains, offering a comprehensive view of the dynamic interactions governing protein states.

Consistency across different systems and mutants reinforced the validity of our model’s predictions. In the Q205L mutant, the rigidity of the Switch II segment prevents the protein from reverting to the inactive state, leading to persistent activation and indicating an enhanced binding capacity to the GTPase domain. This persistent activation is attributed to altered movement patterns between Switch II and other regions, such as the GTPase domain.

In conclusion, our study presents an effective approach for investigating the conformational dynamics of Gαo using a Transformer-based graph neural network. The DMFormer model demonstrated high predictive accuracy and valuable interpretability through integrated gradient analysis. Insights into the persistent activation of the Q205L mutant significantly enhance our understanding of protein activation mechanisms. Our findings have important implications for designing therapeutic interventions targeting protein dynamics and offer a promising avenue for future research in molecular dynamics simulations and protein engineering.

## Acknowledgement

We thank the National Natural Science Foundation of China (22073018, 22377015) for the financial support.

## Supporting Information Available

Code related to this research is available on GitHub.^25^

